# Modulation of host learning in *Aedes aegypti* mosquitoes

**DOI:** 10.1101/172726

**Authors:** Clément Vinauger, Chloé Lahondère, Gabriella H. Wolff, Lauren T. Locke, Jessica E. Liaw, Jay Z. Parrish, Omar S. Akbari, Michael H. Dickinson, Jeffrey A. Riffell

## Abstract

How mosquitoes determine which individuals to bite has important epidemiological consequences. This choice is not random; most mosquitoes specialize in one or a few vertebrate host species, and some individuals in a host population are preferred over others. Here we show that aversive olfactory learning contributes to mosquito preference both between and within host species. Combined electrophysiological and behavioural recordings from tethered flying mosquitoes demonstrated that these odours evoke changes in both behaviour and antennal lobe (AL) neuronal responses. Using electrophysiological and behavioural approaches, and CRISPR gene editing, we demonstrate that dopamine plays a critical role in aversive olfactory learning and modulating odour-evoked responses in AL neurons. Collectively, these results provide the first experimental evidence that olfactory learning in mosquitoes can play an epidemiological role.

## Introduction

Mosquitoes are notorious for their proclivity in host species preferences, and as some of us can attest, certain individuals are preferred over others (*1-3*). In addition, many mosquito species can shift host species when their preferred blood resource is no longer present (*4-6*). Although the abundance of certain hosts often determines mosquito choice (especially if the species is opportunistic), even mosquitoes with a clear host specialization may shift when their preferred host becomes less abundant (*4,5,7*). For example, the generalist mosquito *Culex tarsalis* in California feeds primarily on birds in the summer but on both mammals and birds in the winter (*5,8*). This alteration is linked to fall migration of robins, the mosquitoes’ preferred host. For the highly anthropophilic species *Anopheles gambiae*, in an environment where humans are not readily accessible, >80% of mosquitoes still show an innate preference for human odour, even though the proportion of human feeds is low (<40%)(*4*). This suggests that the mosquitoes have evolved a plastic strategy of feeding on readily available but less preferred hosts.

How do mosquitoes alter their preferences? Although genetic factors may be important (e.g. presence of conserved olfactory receptors to host odours), physiological factors and the mosquitoes’ learning experiences using other blood hosts are likely mechanisms guiding these shifts (*6*). Over the last decade, evidence of olfactory learning in blood-feeding insects has grown for mosquitoes (*9-12*) and kissing bugs (*13-15*). Despite this evidence, the neurophysiological and molecular bases for learning in insect vectors remain unknown, and it is unclear how experience influences host preferences.

For mosquitoes, hosts serve as both prey (source of food, i.e. blood) and predator. The host’s anti-parasitic and defensive behaviours are a major source of mortality for adult female mosquitoes (*16*). Here we take advantage of host defensive behaviours to examine the ability of mosquitoes to learn the association between host odours and aversive stimuli. Our results show that dopamine-mediated olfactory learning is the basis by which mosquitoes aversively learn to shift host preferences, and this strongly modulates antennal lobe (AL) neurons, thereby increasing the mosquito’s ability to discriminate between and learn new hosts.

## Mosquitoes learn to avoid host odours

When encountering a defensive host, mosquitoes are exposed to mechanical perturbations (e.g. swatting, shivering) that can be perceived as negative reinforcement by the insect when paired with other host-related cues such as host odours. Learning the association between host odour and mechanical perturbation would allow mosquitoes to use information gathered during previous host encounters. To determine whether mosquitoes can aversively learn human body odour, 6-day-old mated *Aedes aegypti* females were trained in small individual chambers to associate host-related odorants (conditioned stimulus, CS) with an aversive stimulus consisting of mechanical shocks/vibrations (unconditioned stimulus, US) mimicking host defensive behaviours (Fig. 1A). Twenty-four hours post training, the behavioural response of mosquitoes was assessed in a Y-maze olfactometer in which the insects had to fly upwind and choose between one arm delivering the test odour (i.e. the CS odour) and a control arm carrying only the solvent control (Fig. 1B).

**Figure 1.**
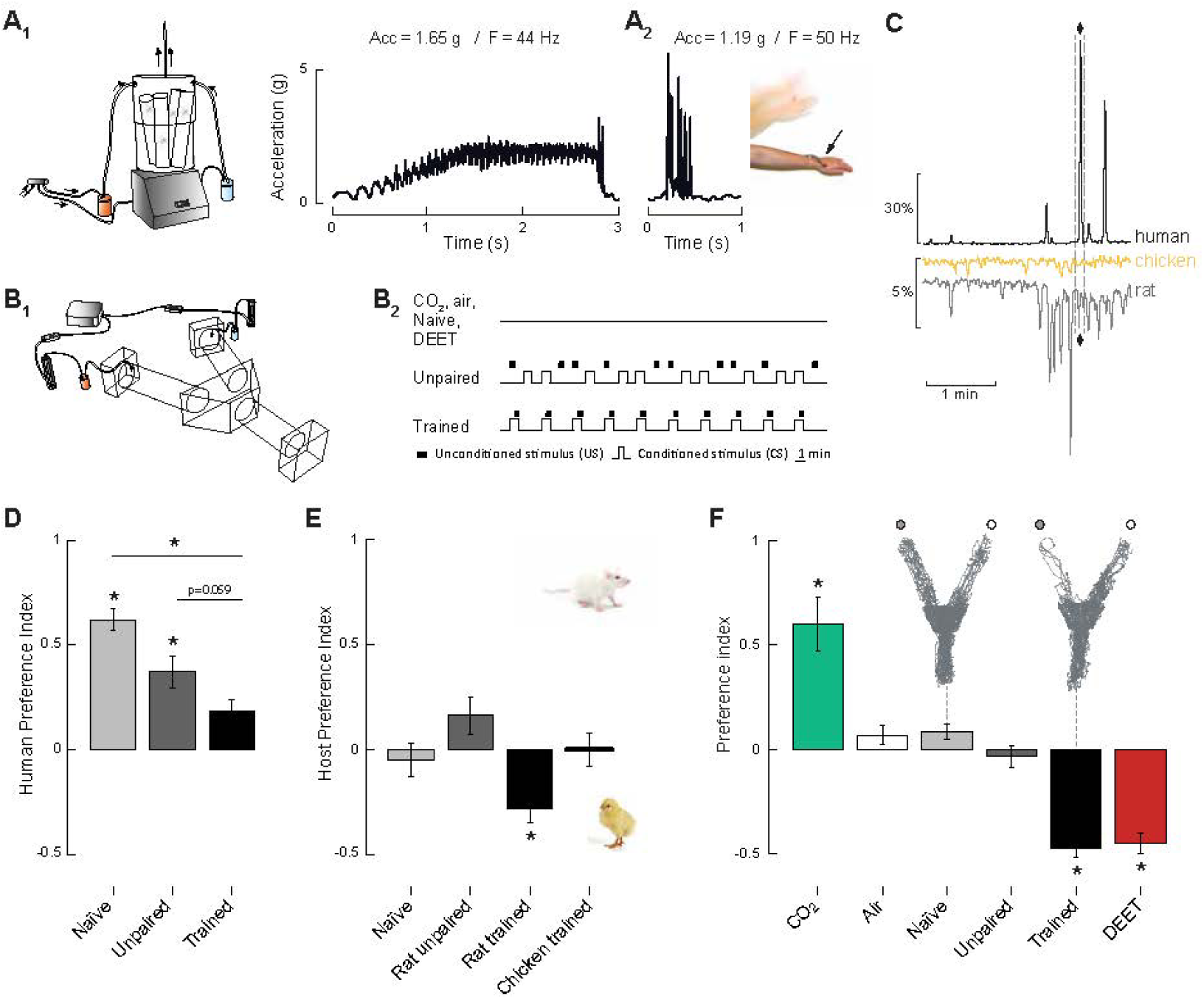
Mosquitoes aversively learn host odours and single odorants. (**A_1_**) *Left*: Aversive training device: mosquitoes are enclosed in individual tubes and stimulated with a mechanical shock from the vortexer and odour (or solvent control) delivered from a scintillation vial. *Right*: Accelerometer recordings from inside the training device and (**A_2_**) from an individual’s arm allowed us to scale the appropriate forces experienced by a mosquito exposed to host defensive behavior. (**B_1_**) Y-maze olfactometer used in behavioral experiments. Mosquitoes are released in the starting chamber, fly upwind, and then have the choice between two arms, each delivering a different odour stimulus. (**B_2_**) Sequences of event delivery [i.e. shock (unconditioned stimulus, US), odour (conditioned stimulus, CS), and inter-trial interval (ITI)] during the experiments. (**C**) Representative GCMS chromatograms of the different host species: human (black, top), chicken (middle, yellow), and rat (grey, bottom). The octenol peak is indicated by the diamond sign. (**D**) Mosquito human host preference represented as a preference index computed from the distribution of insects in the olfactometer. (**E**) Mosquito host preference between the rat and the chicken scents, represented as a preference index. (**F**) Mosquito preference for a CO_2_ positive control (green bar), a DEET negative control (red bar), and octenol (all other bars). Above the naive and trained groups, flight trajectories of individual mosquitoes in response to octenol (grey circle) and a control (white circle). (D-E) Each bar is the mean +/- se from 15-71 mosquitoes; asterisks denote responses that are significantly different from random (binomial test: p<0.05).

*Ae. aegypti* mosquitoes have a strong preference for human hosts (*17,18*). The body odours of individual human subjects (3 males, 3 females) were collected with nylon sleeves (Fig. 1C, S1), and mosquito responses were tested in the Y-maze olfactometer. Whereas naive mosquitoes were strongly attracted to human body odours (Fig. 1D), trained mosquitoes had significantly reduced attraction levels. This reduced attraction shown by trained mosquitoes was not a function of the physiological stress or number of active individuals, but rather was an active decision to avoid the previously experienced odour and fly into the control arm (p>0.05, *t*-test comparisons of flight velocities and activity levels, *n*=24-39; *t*>3.8; for all treatments depicted in Figs. 1, S2). As an important control, we exposed mosquitoes to the CS and US in an unpaired way, thereby preventing the temporal contingency between the stimuli. These mosquitoes, although displaying slightly lower attraction than naive mosquitoes, were still significantly attracted to human odours. Interestingly, not all human subjects elicited the same levels of attraction in naive mosquitoes, and learning performances differed between groups of trained mosquitoes as a function of the individual human body odour used as a CS (Fig. S1).

To test whether associative learning could also affect host selection processes at interspecific levels, rat and chicken body odours were collected using similar nylon sleeves and used in training. The preference of mosquitoes for one of the two host species was tested in the Y-maze olfactometer 24 h after training. In this experiment, one arm delivered the rat odour while the other delivered the chicken odour. Whereas naive mosquitoes and mosquitoes from the unpaired group were equally attracted to the scent of the two host species, mosquitoes trained against the rat odour were significantly more likely to avoid the rat arm and flew preferentially into the arm delivering the chicken odour (Fig. 1E). Conversely, training did not affect mosquito choice when the chicken odour was used as a CS (Fig. 1E). These results mirror those obtained in the triatomine bug, *Rhodnius prolixus,* where bugs successfully learned the association between rat body odours and a mechanical shock but did not learn as well when bird odour was used as a CS (*15*).

The scents emitted by humans and other hosts are complex mixtures of hundreds of odorants, making it difficult to identify which features the mosquitoes might be using to learn the association. We therefore examined the learning capabilities of mosquitoes to single odorants, several of which are emitted from hosts. One that elicited clear learning responses was 1-octen-3-ol (octenol), a common odorant found in the headspace of mammals (*19,20*), but missing in birds (Fig. 1C). We therefore used octenol to more fully explore the ability of mosquitoes to learn the association between the shock and a single host-related odorant. Twenty-four hours after training, mosquitoes remembered the association between the mechanical shock and octenol (Fig. 1F), and their aversive response was comparable to the responses of naive mosquitoes to 40% DEET (N,N-diethyl-meta-toluamide), a concentration corresponding to commercially available doses of this common insect repellent. Again, mosquitoes from the unpaired group did not show learned responses to octenol, clearly demonstrating the associative nature of their learning.

## Aversive learning modifies odour-guided feeding preferences and tethered flight responses

Evidence that learning modifies mosquito olfactory flight preference does not necessarily mean that biting and landing preferences might also be modulated. To examine this, we trained groups of mosquitoes using our aversive learning paradigm (Fig. 1A) and released them into a cage in which they had access to two artificial feeders filled with heparinized bovine blood (37° C); one feeder was scented with octenol while the other was unscented (Fig. 2A). Significantly fewer trained mosquitoes landed on the octenol feeder compared to the control feeder (p<0.0001, binomial test; Fig. 2B). Once they landed, an equal proportion of trained mosquitoes initiated probing on the two feeders (p=0.32, paired Student’s *t*-test, *n*=10; *t*=-1.03; Fig. 2C), although we did observe a tendency for the mosquitoes to feed more on the control feeder than the octenol feeder (24.6 % and 15.6 % of mosquitoes that landed initiated feeding, respectively; p=0.057, binomial test; Fig. S3). By contrast, naive mosquitoes demonstrated no preference in their landing and biting responses to the two feeders (p=0.22, binomial test). The unpaired group showed a slight but significant increase in the proportion of mosquitoes that landed on the scented feeder (p=0.002, binomial test), suggesting that prior exposure to octenol modified their responses in this context. Together, these results suggest that olfactory learning mediates long-(>1 m) and short-range (∼0.1 m) discrimination by the mosquitoes, but once they land, other cues (e.g. heat, water vapour) may partially override these responses (*21,22*).

**Figure 2.**
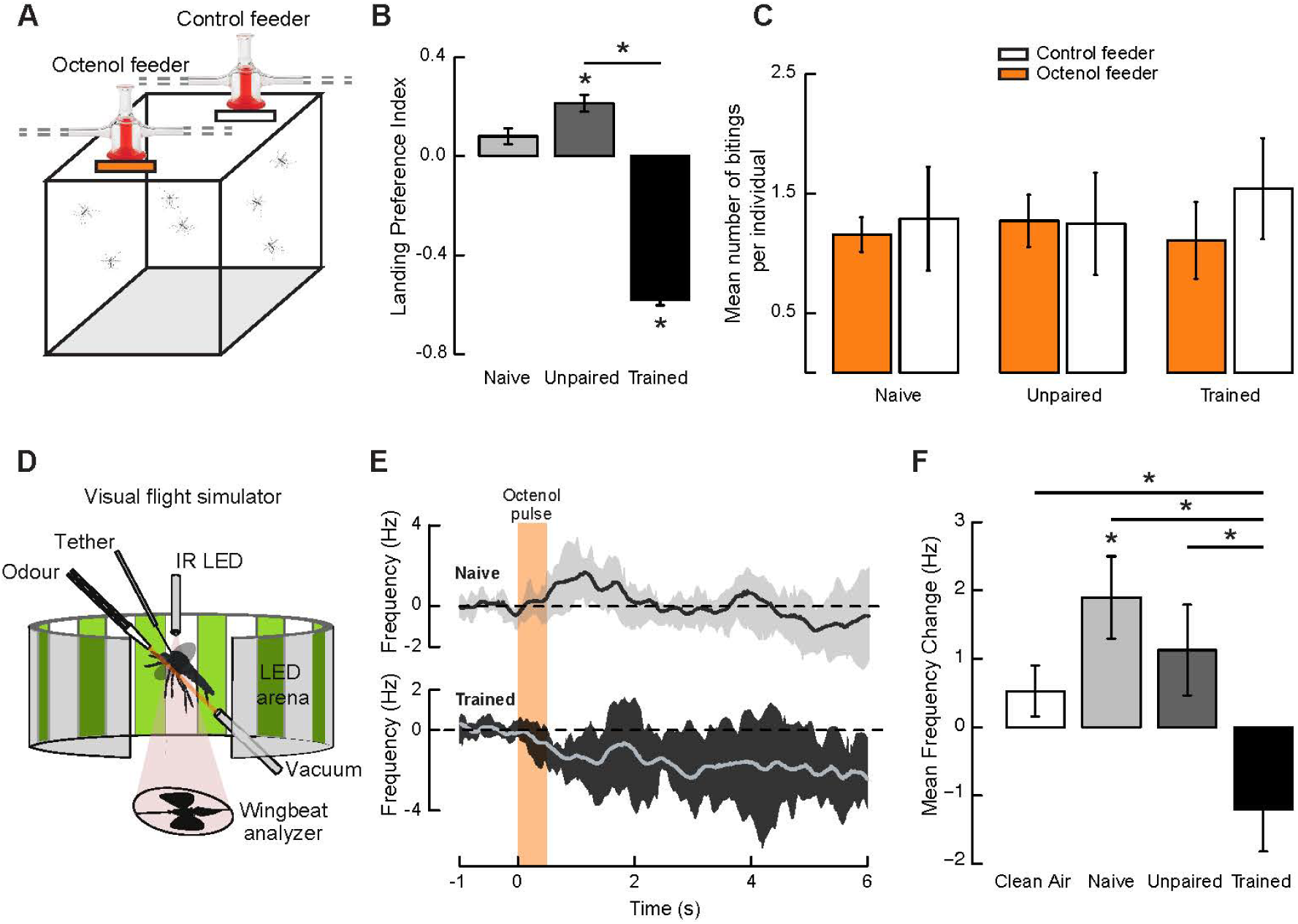
Aversive learning modifies odour-guided feeding preferences and flight responses. (**A**) Experimental setup for testing mosquitoes odour-guided feeding and biting behaviour, each feeder provided heparinized bovine blood and were scented with either octenol or water. (**B**) Mosquito landing preference index for either one of the two artificial feeders, for the naive, unpaired and trained groups. Bars are the mean +/- se, with each bar representing 9-10 groups of 17 responsive female mosquitoes; asterisks denote distributions that are significantly different from random (binomial test: p<0.05). (**C**) Average number of biting per individual on each of the two feeders for the naive, unpaired and trained groups. (**D**) Visual flight simulator (*48,49*) used to record wing kinematics from a tethered mosquito. (**E**) Stimulus-trigger-averaged changes in wingbeat frequency (solid line) in response to a pulse of octenol (light orange bar) for the naive and the trained groups. Shaded areas represent the mean ± the first quartiles. (**F**) frequency response to a pulse of air (white bar) or octenol for the naive (light grey bar), unpaired (dark grey bar) and trained (black bar) groups. Each bar is the mean +/- se of 16-23 responsive female mosquitoes; asterisks denote significant responses compared to zero when located above bars, or between groups when located above horizontal lines (p<0.05, Student’s *t*-test, *t*>1.57).

To better understand how learning modulates flight responses and to determine whether mosquitoes fly while tethered (thereby allowing simultaneous behavioural analysis and electrophysiological recordings from the AL), we positioned mosquitoes in the centre of a virtual LED arena where they were tethered by the thorax and maintained in a laminar airflow (Fig. 2D). An infrared (IR) light and a two-sided IR sensor allowed real-time measurements of the mosquitoes’ wingstroke frequency, amplitude, and turning tendency. Results showed that whereas naive and unpaired mosquitoes exhibited a frequency increase in response to a brief octenol pulse, trained mosquitoes significantly decreased their flight frequency in response to the same stimulus (p=0.013, Student’s *t*-test, *n*=34; *t*=2.67; Figs. 2E,F; S4).

## Dopamine is critical for aversive learning

Classical insect models for studying learning and memory have shown that dopamine is a key neuromodulator involved in aversive learning (*23-26*). To test whether dopamine is also implicated in aversive learning in mosquitoes, we used several ways to manipulate dopamine receptors, including dopamine receptor antagonist injections (Fig. 3A, top-left), gene knock-down via RNAi (Fig. 3A, top-centre) and CRISPR/Cas9 gene-editing methods (Fig. 3A, top-right). After aversive training to octenol, mosquitoes were tested in the Y-olfactometer (Fig. 1B), allowing us to quantify their flight velocities and behavioural preferences. First, adult female mosquitoes that received dopamine receptor antagonist injections showed significant deficits in their learning abilities compared to uninjected and saline-injected mosquitoes, which showed robust learning responses (Fig. 3B). Similarly, female mosquitoes that were injected with dsRNA targeting the *DOP1* gene and CRISPR mutants with a 6-amino acid deletion of the *DOP1* receptor (Fig. S5) showed significant learning deficits compared to the uninjected, non-target dsRNA injected and saline-injected control groups (p<0.05, binomial test compared to control groups; Figs. 3B, S6). There were no significant differences in the responses of mosquitoes in treatment groups in which the dopamine receptor was manipulated (i.e. antagonist injected, dsRNA injected, CRISPR edited; p>0.64, binomial test). To evaluate the effects of dopamine receptor manipulation on flight responses, we quantified the mosquito flight trajectories in the olfactometer. Results showed that there was no significant difference in flight velocity between dopamine-impaired treatment groups or between those groups and the saline-injected and uninjected controls (p>0.05, Student’s *t*-test, pairwise comparisons Holm p-value adjustment, *n*=17-29; *t*<2.03; Fig. S2), suggesting that dopamine receptor manipulation did not affect mosquito flight-motor responses. However, it is worth noting that dsRNA-injected mosquitoes and *DOP1* mutants were significantly less aroused to the odours than the other treatment groups (p<0.05, binomial test; Fig. S2). Nonetheless, when these dopamine-impaired mosquitoes were tested against CO_2_ or human host odours, they all showed significant attraction (p<0.05, binomial test; Fig. 3C,D), revealing that manipulating the dopamine receptors impaired their ability to learn aversive information but did not affect their innate olfactory behaviour.

**Figure 3.**
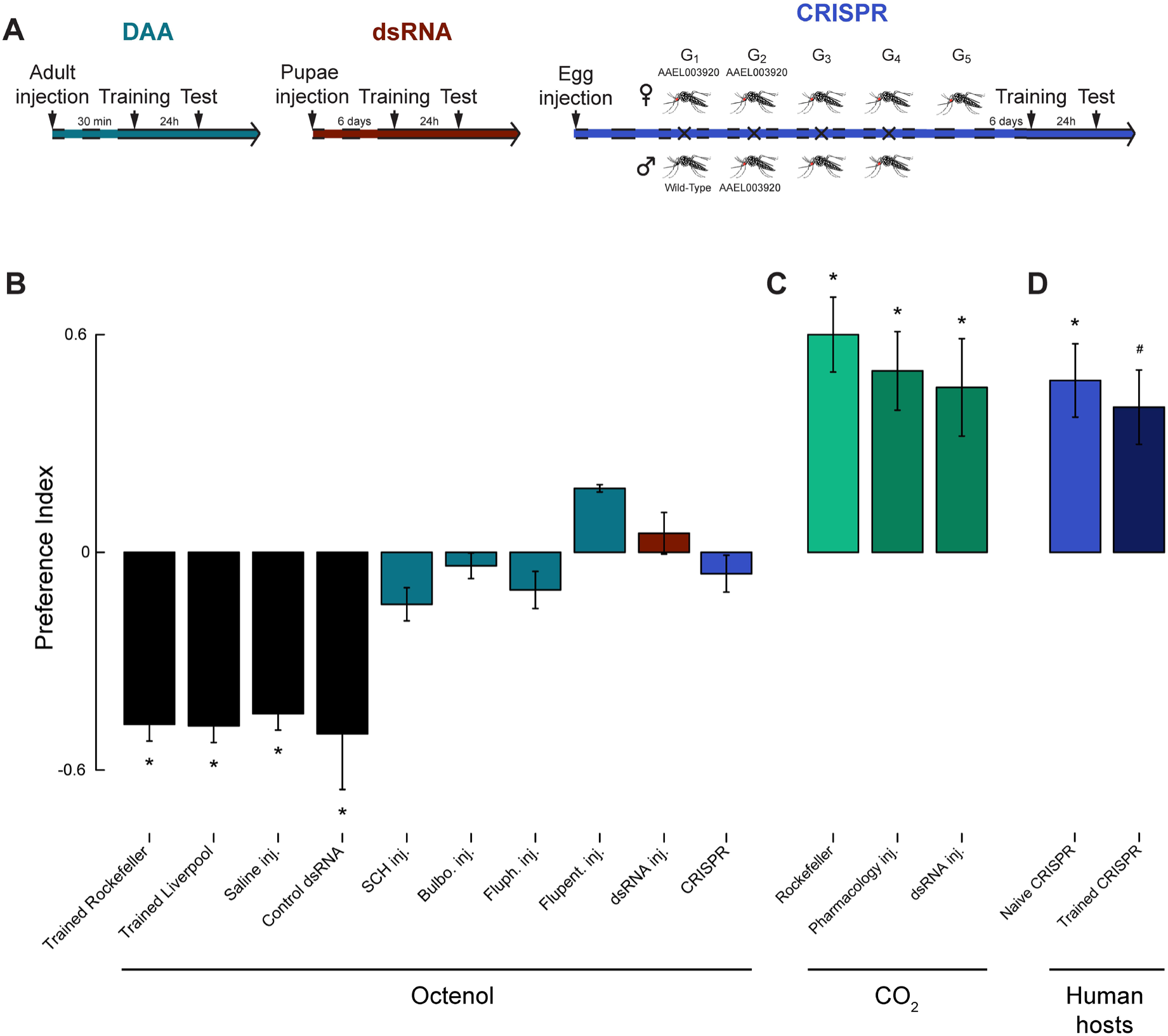
Dopamine is involved in mosquito learning. (**A**) *Left*: Dopamine receptor antagonists (DAA) (SCH-23390, bulbocapnine, flupentixol and fluphenazine) were injected in the thorax of 6-day old female mosquitoes that were trained 30min post injection and tested 24 hrs later. *Centre*: *DOP1* and control dsRNA were injected in 1-day old pupae, and after 6-d post emergence mosquitoes were trained and tested. *Right:* CRISPR/Cas9 constructs were injected in embryos. Mutants were screened and selected by sequencing for 5-8 generations before being trained at 6 days old. (**B**) Mosquito choice in the olfactometer represented as a preference index. Trained mosquitoes from the Rockefeller, Liverpool strain, saline injected and dsRNA injected Rockefeller lines were not significantly different in their learning performances (p>0.05, binomial test; black bars). By contrast, mosquitoes injected with dopamine receptor antagonists (blue-green bars), dsRNA-injected (red bar), and CRISPR mosquitoes (mauve bar) showed no learning. Mosquitoes injected with dopamine receptor antagonists (SCH-23398, 10^-6^M) or dsRNA, as well as CRISPR mosquitoes were still responding to positive controls such as CO_2_ (**C**) or host odours (**D**). When human scents were used during training, CRISPR mosquitoes showed no learning (p=0.79, binomial test). Each bar (mean +/- se) representing 11-29 responsive female mosquitoes; asterisks indicate distributions that are significantly different from random (p<0.05, binomial test); # indicates p<0.06 when the response of the trained CRISPR was compared to chance.

Given the inability to learn octenol by the *DOP1* mutants, how might they respond to human scent that contains hundreds of volatiles that are highly attractive to mosquitoes? Results showed that naive *DOP1* mutants were significantly attracted to the scent of human hosts that were also attractive to wild type mosquitoes (male#1, #2 and female #1; p<0.05, binomial test; Figs. 3D, S1). Trained *DOP1* mutants failed to learn the association between the shock and human odours, exhibiting similar behavioural responses to the naive mosquitoes (p=0.79 when compared to the naïve CRISPR tested against human odours, binomial test; Fig. 3D). Moreover, responses by the trained *DOP1* mutants contrasts those of the trained wild type mosquitoes, which showed learned aversive responses to those same hosts (Fig. 1D).

## Odour stimuli are learned and represented distinctly in the mosquito brain

Given the differences in mosquito olfactory preferences between human and vertebrate hosts and previous work showing that only certain odour stimuli can be learned (*12*), we next examined how mosquitoes learn different odorants and how odour stimuli are represented in the brain. Twenty-four hours after training, behavioural responses showed that mosquitoes did not learn all odorants equally. For example, whereas responses to nonanol were not influenced by aversive training, those to octenol showed learned aversive responses and L-(+)-lactic acid caused significant attraction (Figs. 1F, 4A). To evaluate how different host- and plant-associated odorants are represented in the mosquito brain, we performed extracellular recordings of projection neurons (PNs) and local interneurons (LNs) in the antennal lobe (AL), simultaneous with behavioural recordings (Fig. 4B). The extracellular recording method did not allow us to distinguish between PNs and LNs, but it did provide stable recordings (>1 h) of multiple neural units (Fig. S7) while allowing us to simultaneously quantify odour-evoked changes in wingbeat amplitudes. Whereas the mineral oil (no odour) control elicited no change in behavioural and neural responses, stimulation with octenol and ammonia elicited strong firing rate responses in single units (Fig. 4C). Interestingly, whereas ammonia elicited a one to two seconds change in wingstroke activity, stimulation with octenol elicited much longer behavioural responses that lasted many seconds beyond the duration of the stimulus (400 ms) (Fig. 4C). Examining single-unit responses across the odour panel, we found that the majority of units (∼65 %) showed strong odour-evoked responses, with the remaining units showing no significant change in activity (Figs. 4D, S8). Moreover, some units (19 %) were broadly responsive to different odorants, including units that were responsive to aromatics (e.g. benzaldehyde) and aliphatic compounds (e.g. octenol), as well as monoterpenes (e.g. D-limonene) (Fig. S8). By contrast, others (27 %) were more narrowly tuned, including units that only responded to one chemical class. In these experiments, hexanol, hexanal, butyric acid, cresol, DEET, ammonia, and breath evoked behavioural responses that were significantly higher than observed for the control (p<0.05, pairwise Student’s *t*-tests with Holm correction for multiple comparisons, *n*=10-16; *t*>2.38). Interestingly, the behavioural state (i.e. flying or non-flying) had a significant effect for units that showed suppressed firing activity when stimulated with an odour (p<0.01, Kruskal-Wallis rank sum test, χ^2^=6.95) but not for units that showed excitatory responses (p=0.51, Kruskal-Wallis rank sum test, χ^2^=0.44). It is also worth noting that the spontaneous activity of units was slightly (but not significantly) higher when the mosquitoes were flying (p=0.083, Kruskal-Wallis rank sum test, χ^2^=3.01).

**Figure 4.**
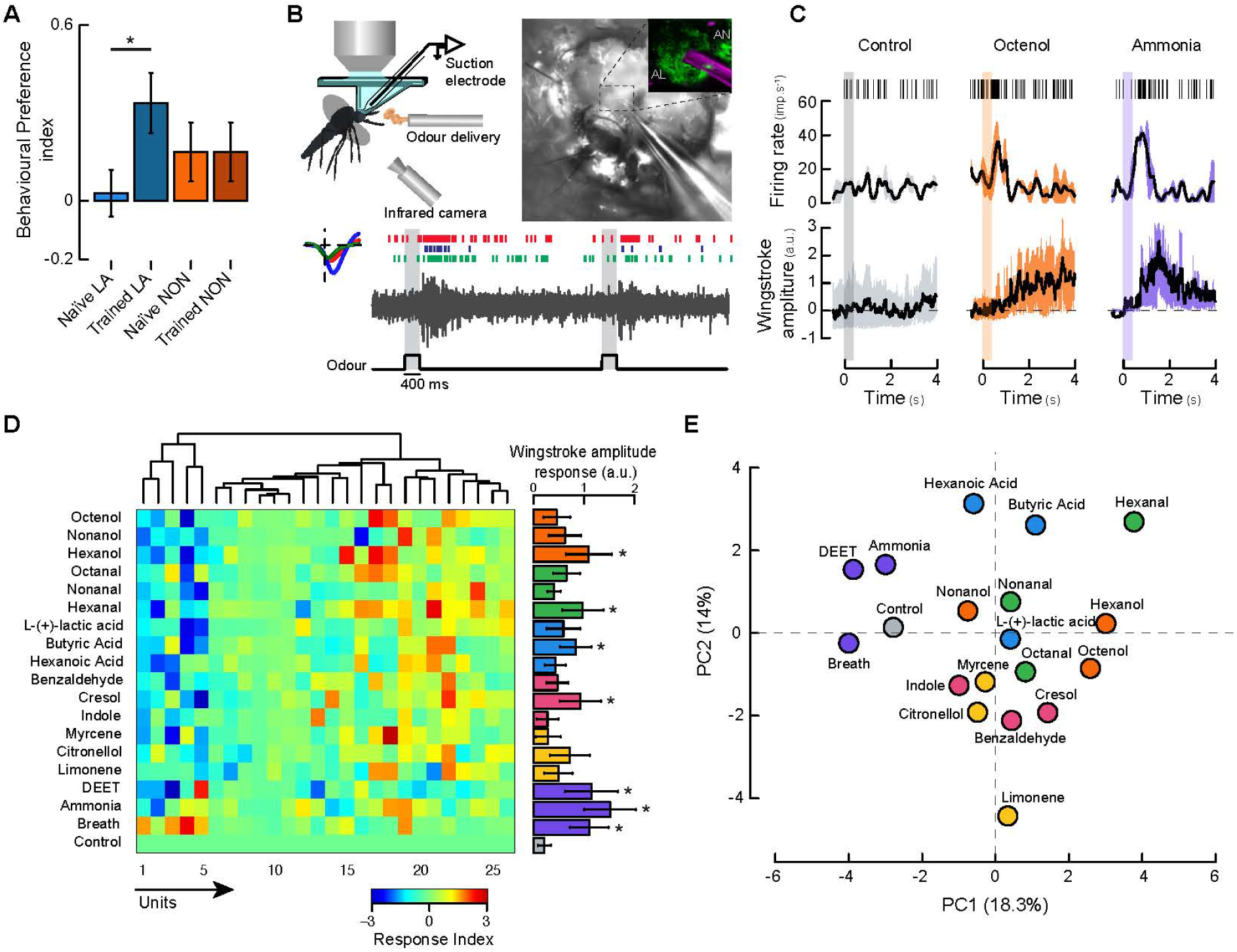
Odour stimuli are learned and represented differentially in the mosquito brain. (**A**) Mosquito preference index (PI) for L-(+)-lactic acid (LA; blue bars) and 1-nonanol (NON, orange bars), tested in the olfactometer. Each bar is the mean +/- se from 21-39 responsive female mosquitoes; asterisks denote p<0.05 (binomial test). (**B**) *Top,left*: Electrophysiological preparation for simultaneous flight behaviour and suction electrode recording from the mosquito antennal lobe (AL), which receives olfactory input from the antenna and maxillary palps. *Top,right*: Picture of the suction electrode inserted in the right AL of a mosquito. *Inset*: representative electrode position (5µm tip diameter, purple) relative to the AL (green) and antennal nerve (AN). *Bottom*: Representative raw recording and raster plot showing the responses of three units after the delivery of 400 ms pulses of octenol (grey bar). (**C**) *Top*: Raster plots and peri-event histograms of the mean (± variance) responses of an isolated unit from the suction electrode recordings. *Bottom*: stimulus trigger-averaged responses in wingstroke amplitude (± first quartiles) to olfactory stimulation. Vertical shaded bars represents the odour stimulus: clean air (grey), octenol (orange) and ammonia (purple). (**D**) *Left*: Neural ensemble response to the odour panel (rows 1-19), plotted as a colour-coded response matrix across neural units (columns) (*n*=8 preparations). *Right*: normalized change in mean wingstroke amplitude (a.u. ± se) in response to each odour of the panel. Asterisks denote responses that are significantly different from the control (Student’s *t*-test: *n*=10-16; *t*>2.38; p<0.05). (**E**) Principal components analysis of the ensemble responses. a-e: color fills are indicative of the chemical class of the odorant (orange: alcohols, green: aldehydes, blue: carboxylic acids, pink: aromatic and phenolic compounds, yellow: monoterpenes, purple: other compounds, grey: mineral oil control).

At the neural population level, ensemble responses showed distinct clustering in the multivariate (Principal Component Analysis) space based on the type and chemical class of the olfactory stimuli (p<0.001, Kruskal-Wallis rank sum test, χ^2^=12.19; Fig 4E). For example, monoterpenes and aromatics like D-limonene, *β*-myrcene, benzaldehyde, and cresol occupied a distinct region of the olfactory space relative to the aliphatic acids, alcohols, and aldehydes. By contrast, odour stimuli that evoked strong responses across the ensemble (DEET, ammonia, and breath) were grouped together and were significantly different from the other odorants (p<0.001, Kruskal-Wallis rank sum test, χ^2^=11.57), demonstrating that the AL neural ensemble can generalize among and discriminate between olfactory stimuli.

## Dopamine selectively modulates AL neurons

To examine how dopamine modulates the processing of olfactory information, we first used immunohistochemistry to examine dopaminergic innervation (via tyrosine hydroxylase, a dopamine precursor) in the mosquito brain. We found extensive dopaminergic innervation across the brain but particularly concentrated in the ALs and lateral protocerebrum, including the mushroom bodies (Fig 5A), which are centres that mediate olfactory learning and memory in insects (*27,28*). Dopaminergic innervation is heterogeneous in the AL, with some glomeruli being more innervated than others, including the MD2 glomerulus that receives input from the octenol-sensitive aB2 neuron in the maxillary palp (Fig. S9). Antisera against the D1-like dopamine receptor *DOP1* reveal staining of cell bodies around the ALs, as well as enrichment in the lateral protocerebrum surrounding the mushroom bodies (Figs. 5A, S9B). We therefore sought to determine the effects of dopamine on odour-evoked responses of mosquitoes AL neurons.

**Figure 5.**
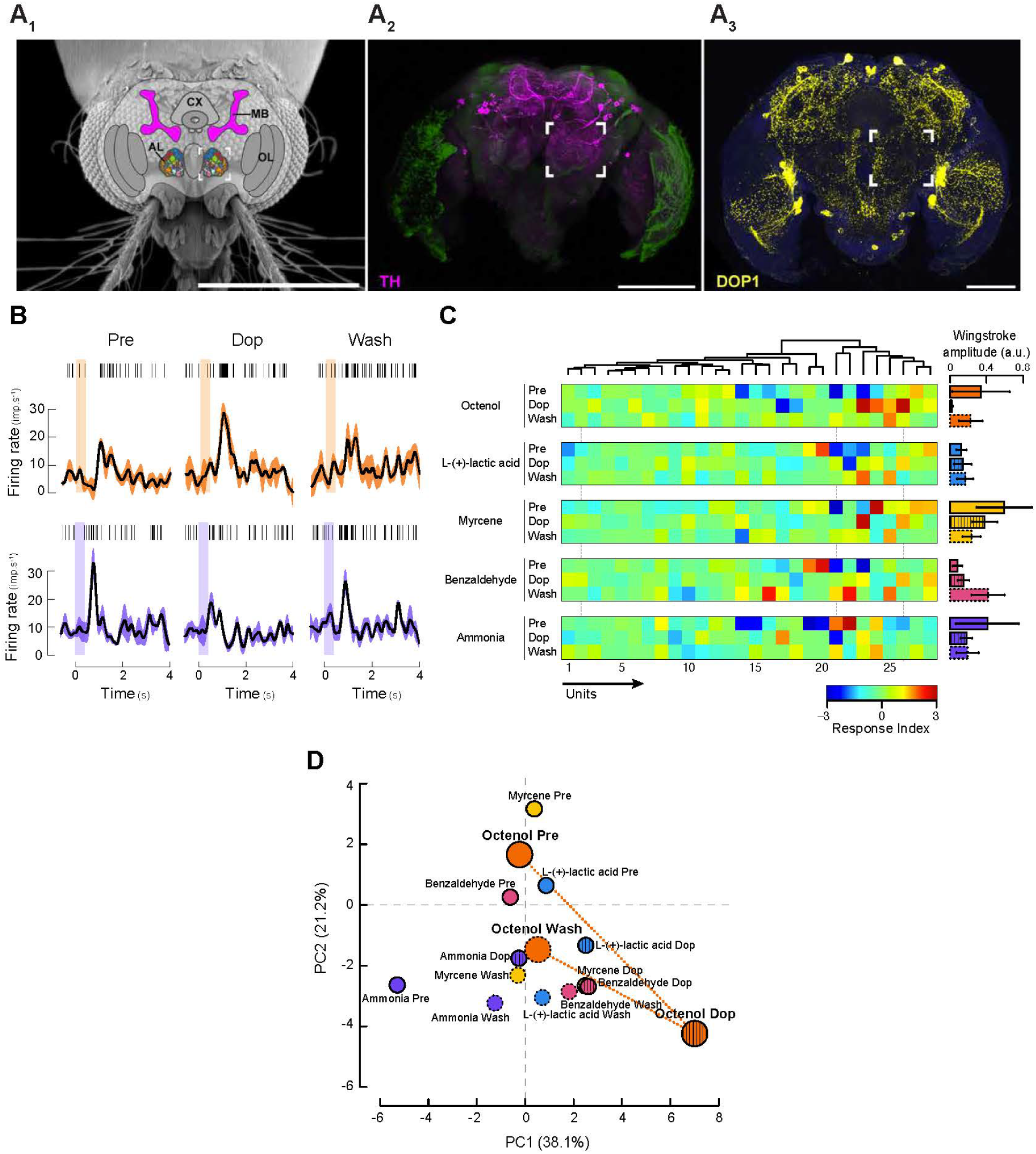
Dopamine selectively modulates antennal lobe neurons. (**A_1_**) Schematic of the *Ae. aegypti* brain superimposed on a scanned electron microscope image (*50*). Highlighted regions include the AL (multicolored to represent individual glomeruli that receive input from olfactory receptor neurons), and the mushroom bodies (MB), implicated in learning and memory. The open box around the AL is used to indicate the corresponding location in **A_2_** and **A_3_**. CX: central complex; OL: optic lobes. Scale bar: 500 µm. (**A_2_**) Confocal micrograph of a whole *Ae. aegypti* brain stained with antibodies against tyrosine hydroxylase (magenta) shows immunoreactivity concentrated in the lateral protocerebrum and AL. Background fluorescence in green. Scale bar: 100 µm. (**A_3_**) A 60 µm section of *Ae. aegypti* brain stained with antibodies against the mosquito dopamine-1 receptor-1 *DOP1* (yellow) shows these receptors enriched in the lateral protocerebrum around the MB as well as localized around the AL. Background fluorescence in blue. Scale bar: 100 µm. (**B**) *Top*: Raster plots and peri-event histograms of the mean (± variance) responses of an isolated unit from the suction electrode recordings. *Bottom*: stimulus trigger-averaged responses in wingstroke amplitude (± first quartiles) to olfactory stimulation. Vertical shaded bars represent the odour stimulus: octenol (orange) and ammonia (purple). Each column corresponds to the responses before (Pre), during (Dop) and after (Wash) dopamine application. (**C**) *Left*: Neural ensemble response to a subset of 5 odorants (octenol, lactic acid, myrcene, benzaldehyde, ammonia) before (Pre), during (Dop) and after (Wash) dopamine application. Responses are plotted as a color-coded response matrix across the neural units (columns). Vertical dashed lines indicate examples of units for which the response either does not change (unit 2), decreases (unit 21 for ammonia) or increases (unit 26 for all odours except ammonia) during dopamine application. *Right*: normalized mean wingstroke amplitude change (a.u.) in response to each odour of the panel, before (open bars), during (hatched bars) and after (dashed bars) dopamine application. Bars are the mean ± se. (**D**) Principal components analysis of the ensemble responses. Borders and colour fills are indicative of the odorant (orange: octenol, blue: lactic acids, pink: benzaldehyde, yellow: myrcene, purple: ammonia) and of the treatment (solid line: Pre, hatchings: Dop, dashed lines: Wash).

To test for the neuromodulatory role of dopamine in mosquitoes, we simultaneously recorded the electrophysiological and behavioural responses evoked by a sub-panel of odorants comprised of octenol, L-(+)-lactic acid, *β*-myrcene, benzaldehyde, and ammonia before, during, and after superfusion of dopamine (1 µM) over the brain. Dopamine application increased odour-evoked firing rate responses (Fig. 5B,C) in 69.6 % of responsive AL units, decreased responses in 21.7% of units, and had no effect in 8.7% of units. Dopamine also increased the sensitivity of ∼17% of the recorded units, leading to a higher number of cells responding to olfactory stimuli. These effects could be washed out in approximately 50% of units and, in contrast to preparations that were superfused with dopamine, additional control experiments with mosquitoes that were continuously superfused with saline showed no change in spontaneous responses (p>0.05, pairwise comparisons using *t*-tests with pooled SD, *t*<1.52; Fig. S10). Moreover, at the level of the neural ensemble, odorant representation significantly changed during dopamine application compared to the pre- and wash-phases of the experiment (p<0.05, Kruskal-Wallis rank sum test, χ^2^=6.17) causing stimuli —in particular, octenol—to become more separated in the olfactory space (Fig. 5D). Interestingly, the degree of modulation was not the same for all odorants, suggesting that the observed heterogeneity in dopaminergic innervation of glomeruli may be functionally linked to glomerular response modulation (Fig. 5A,D).

## Discussion

Heterogeneity in mosquito biting and consequently host infection plays an important role in the spread of vector-borne disease (*29,30*), and previous studies have documented interindividual differences in attractiveness to mosquitoes (*2*), as well as an ability for mosquitoes to shift species when their preferred host is no longer available (*4,31*). Despite these studies, the processes mediating these mosquito behaviours have remained unclear (*32*). Here we show that learning can contribute to these host shifts, and that their direction seems to be driven by the composition of the host odour. In particular, our results show that human individuals that are highly attractive to mosquitoes are the ones that mosquitoes can learn to avoid. Mosquito learning may thus partially explain host preference heterogeneity and flexibility, and it may also elucidate which olfactory channels mediate these changes.

Here in this study we employed an integrative approach to demonstrate that mosquito learning can influence both specificity for individual hosts and their flexibility in olfactory preferences. The ability of mosquitoes to aversively learn depended on odorant type, for instance, L-(+)-lactic acid, an odorant emitted by hosts, could be learned in an appetitive but not aversive context (*12*), whereas octenol—another odorant emitted by both plants (*33*) and blood hosts (*19,20*)—could be appetitively and aversively learned, suggesting that certain odorants may be encoded by specific olfactory channels that allow rapid learning of attractive or defensive hosts or other important odour sources (e.g. carbohydrates). Our electrophysiological recordings revealed that the AL represented the odorants by chemical class and activity level, and dopamine—a critical neuromodulator involved in learning and arousal (*34*)—further increased the separation of those odorants in the AL encoding space. *DOP1* is critical for mediating this plasticity in AL responses and learning abilities, with CRISPR mutants for this receptor showing an inability to learn. Host defensive behaviour is a major source of mortality for mosquitoes, with hosts operating as both predator and prey. Thus, the ability to learn may have strong fitness consequences for the mosquitoes. CRISPR has been highlighted an important tool in the fight against vector-borne disease (*35,36*). Notably, these mutants have allowed us to target the dopaminergic pathway and impair mosquitoes’ ability to use their experience to fine-tune their responses to host signals. Identifying the mechanisms and pathways enabling flexibility in mosquito behaviour may provide tools for more effective mosquito control.

## Material and Methods

### Mosquitoes rearing and colony maintenance

Multiple strains of *Aedes aegypti* mosquitoes were used for the experiments: Rockefeller (ROCK), Liverpool (LVP-IB12) and CRISPR transgenic line from the Liverpool strain. Mosquitoes were maintained in a climatic chamber at 25±1°C, 60±10% relative humidity (RH) and under a 12-12h light-dark cycle. Mosquitoes were fed weekly using an artificial feeder (D.E. Lillie Glassblowers, Atlanta, GA, USA; 2.5 cm internal diameter) supplied with heparinized bovine blood (Lampire Biological Laboratories, Pipersville, PA, USA) and heated at 37° C using a water-bath circulation (HAAKE A10 and SC100, Thermo Scientific, Waltham, MA, USA). Cotton balls soaked with 10% sucrose were continuously provided to the mosquitoes. Eggs were hatched in deionized water that contained powdered fish food (Hikari Tropic 382 First Bites - Petco, San Diego, CA, USA), and larvae were cultured and maintained in trays containing deionized water and the fish food. For the experiments, groups of 100 to 120 pupae (both males and females) of the same age were isolated in individual containers and maintained exclusively on 10% sucrose after emergence (i.e. no blood-feeding). Six-day-old female mosquitoes were individually isolated in 15 mL conical Falcon^TM^ tubes (Thermo Fisher Scientific, Pittsburgh, PA, USA) covered by a piece of fine mesh that permitted odour stimulation during training. Experiments were conducted when the mosquitoes were the most active and responsive to host related cues: 2 hrs before their subjective night (*12,37*).

### Host odour collection and GCMS analysis

Host body odours were collected using nylon sleeves (Ililily Inc., Irvine, CA, USA) that were worn for 3.5 hrs. For human scent collection, volunteers of various ethnic backgrounds (3 females, 3 males, aged from 23 to 43 years old), wore one nylon sleeve around the ankle and one nylon sleeve around the arm. Both sleeves were used simultaneously to either train or test mosquitoes. Volunteers used fragrance-free detergents and soaps to prevent bias in mosquito behaviour. In addition, we also collected headspace volatiles from adult human volunteers as previously described (*38*) by wrapping a volunteer’s arm in aluminum and piercing the aluminum with a 75um CAR/PDMS SPME fiber (57344-U; Supelco, Bellefonte PA USA). Human scent protocols were reviewed and approved by the University of Washington Institutional Review Board, and all human volunteers gave their informed consent to participate in the research. Scent from rats and chicken hatchlings (from <2 years old male rats and 10-day-old chicken hatchlings; both approximately the same mass) were collected by placing a nylon sleeve around the abdomen for 3.5 hrs (IACUC Protocol # 4385-01). To discriminate between endogenous and exogenous volatiles, controls were performed by keeping clean nylon sleeves in clean, unoccupied rearing containers for the same duration as for the odour collection procedure. Host odours were collected by either the SPME method or by dynamic sorption. The latter method involved enclosing the nylon socks in a nylon oven bag (Reynolds Kitchens, USA). Air was withdrawn from the bag via a diaphragm vacuum pump (400-1901, Barnant Co., Barrington, IL, USA) and passed through a headspace trap comprised of a Pasteur pipette with 50 mg of Porapak™ powder Q 80-100 mesh (Waters Corporation, Milford, MA, USA) packed between two plugs of glass wool (Restek, Belfonte, PA, USA); air was returned to the bag through a charcoal-filter. Headspace collections lasted for 24 hrs. Volatiles were eluted from the traps with 600 μL of 99% purity hexane (Sigma Aldrich, St. Louis, MO, USA), and samples were stored in 2 mL amber borosilicate vials (VWR, Radnor, PA, USA) with Teflon-lined caps (VWR, Radnor, PA, USA) at −80°C until they were run on a Gas Chromatograph coupled to a Mass Spectrometer (GCMS). Fibers were exposed to host volatiles for 1 hr before being run on the GCMS.

Liquid samples were injected (or SPME fibers were exposed) into an Agilent 7890A gas chromatograph (GCMS) with a 5975C Network Mass Selective Detector (Agilent Technologies, Palo Alto, CA, USA). A DB-5 GC column (J&W Scientific, Folsom, CA, USA; 30 m, 0.25 mm, 0.25 μm) was used, and helium was used as the carrier gas at a constant flow of 1 cc.min−1. The oven temperature was 45° C for 3.75 min, followed by a heating gradient of 10 degrees.min^-1^ to 250° C, which was then held isothermally for 10 min. Chromatogram peaks were manually integrated using the ChemStation software (Agilent Technologies), tentatively identified by the NIST library before verification using Kovats Indices and synthetic standards.

### Mosquito training protocol and control groups

A total of 2258 individual female mosquitoes were used in the behavioral experiments. Before each training session, individual mosquitoes were allowed to acclimate for 1 min in the absence of stimulation, except for the delivery of a clean air at 30 cm.s^-1^, room temperature (23° C) and relative humidity (50%). Mosquitoes were then simultaneously exposed to the olfactory stimulus (e.g., octenol at 140 mM; equivalent to the concentrations used in other mosquito training experiments (*12*)) and a mechanical shock that was delivered for 30 sec by a vortexer (Thermo Fisher Scientific, Waltham, MA, USA) at 1.65 g at 44 Hz. Forces were scaled to host defensive behaviours that occur when a human slaps his/her arm to drive off biting mosquitoes (Fig. 1A) as well as exposing mosquitoes to a strong mechanical perturbation without damaging their wings or causing apparent physiological and/or physical damage. Mosquitoes were exposed to ten training trials, each separated by a 2 min interval. During this inter-trial interval (ITI), mosquitoes were maintained in the same experimental room and exposed to a filtered air flow. A vacuum line was used throughout the training session to remove environmental contaminants and olfactory stimuli from the container during the ITI. After conditioning, mosquitoes were placed in a humidified climatic chamber (25° C; 60% RH; 12-12 h L:D) and tested in the Y-olfactometer 24 hrs post-training. Two control groups were used to test for the effects of aversive learning: a “naive” untrained group; and an “unpaired” group. The “unpaired” group controlled for the associative nature of the learning, by exposing mosquitoes to the odour and the mechanical shock in a pseudo-random, unpaired sequence, i.e. in the absence of temporal contingency (*39*). Each of the control groups was tested 24 hrs later.

### Behavioral testing in the olfactometer

We used a custom-made, Plexiglas® Y-maze olfactometer to evaluate and compare mosquito responses to different odour stimuli, as previously described (*12*)(Fig. 1B). Briefly, the olfactometer comprised of a starting chamber, allowing mosquito release, an entry tube (30 cm long, 10 cm diameter) connected to a central box where two “choice” arms were attached (both 39 cm long, and 10 cm diameter). Charcoal filtered air entered as a uniform laminar flow at 20 cm.sec^-1^ into the arms of the olfactometer (Fig. 1B). Odour stimuli were delivered to each choice arm via teflon® tubing connected to one of two 20mL scintillation vials containing either the tested odour or the control solution (mineral oil) (Fig. 1B). Each line was connected to the corresponding choice arm of the olfactometer and placed centrally in the olfactometer arm. All the olfactometer experiments were conducted in a well-ventilated climatic chamber (Environmental Structures, Colorado Springs, CO, USA) at 25°C and 50% RH. After each experiment, the olfactometer, tubing and vials were cleaned up with water followed by 70% and then 100% ethanol to avoid any contamination between experiments. Finally, to avoid any biases, the side of the stimulus and control arms was randomized daily.

Testing sessions began when one single mosquito was placed in the starting chamber. The mosquito then flew along the entry tube and, at the central chamber, could choose to enter one of the olfactometer arms, one emitting the trained stimulus and the other the “clean air” (solvent only) control (*12*). We considered the first choice made by mosquitoes when they crossed the entry of an arm. Mosquitoes that did not choose or did not leave the starting chamber were considered as not responsive and discarded from the preference analyses. Overall, 68.5% of the females were motivated to leave the starting chamber of the olfactometer and choose between the two choice arms. In addition, four treatments were used to ensure that contamination did not occur in the olfactometer and to test mosquitos’ responses to innately attractive or aversive stimuli. Untrained “naive” mosquitoes were placed in the olfactometer and exposed to either: (1) two clean air currents (neutral control); (2) a clean air stream versus CO_2_ (positive control, [CO_2_] = 2300 ppm above ambient level) (*40*); (3) a clean air stream versus 40% DEET (an innately aversive control); or (4) a clean air versus octenol (i.e. naive control). Mosquito trajectories were captured with a video camera (Model C615, Logitech, Newark, CA, USA) (Figs. 1F, S2) and mosquito flight speeds were calculated for each individual. See supplementary information for details on data analysis and statistical tests.

### Behavioral testing with the artificial feeder

In order to test whether mosquitoes could use learned information in the context of blood-feeding, groups of 17 female mosquitoes were released in a cage (30.5×30.5×30.5, Bioquip^Ⓡ^, Rancho Dominguez, CA, USA) on top of which two artificial feeders containing heparinized bovine blood, warmed up to 37° C, were positioned. One feeder was treated with the CS odour (pipetted onto a Kimwipe (Kimberly-Clark professionals, Roswell, GA, USA) surrounding the feeder), while the control feeder (odourless) was treated with the solvent only (i.e. MilliQ water). Two video cameras (Model C615, Logitech, Newark, CA, USA) were used to record mosquitoes’ activity at each feeder over the course of the experiment (25 min duration) (Fig. 2A) and the total number of landing, piercing and feeding events was counted for each feeder. The position of the feeder associated with the CS odour was randomized in order to avoid any potential spatial bias. Tethered flight experiments are described in the supplementary information.

### Interrogation of dopamine pathways in the mosquito brain

To evaluate the impact of dopamine on mosquito olfactory learning, we used three different approaches: 1) dopamine receptor antagonist injections; 2) knockdown of *DOP1* using RNA interference and 3) modification of *DOP1* using the CRISPR/Cas9 method (see supplementary information for details).

#### dsRNA synthesis, precipitation and injection

Double-stranded RNA (dsRNA) of *DOP1* and *Drosophila* nautilus (non-targeting control, #M68897) genes were synthesized by *in vitro* transcription using the MEGAscript® RNAi kit (ThermoFisher Scientific, Waltham, MA, USA - AM1626) following the manufacturer’s recommendations (see supplementary information for DNA template preparation details). The integrity of the products was assessed by agarose gel electrophoresis (0.8%) to ensure that the fragments were of the proper size and not degraded. After synthesis, the dsRNA was precipitated using sodium acetate and ethanol and resuspended in nuclease free water (ThermoFisher Scientific, Waltham, MA, USA). The concentration and integrity of the dsRNA were determined by spectrometry (NanoDrop 2000c, Thermo Scientific, Wilmington, DE, USA) and electrophoresis. The dsRNA was then kept at −80°C until the injections were performed. Before the injection, the dsRNA was thawed and diluted in water to the desired concentration. Injections were performed using a pulled borosilicate pipette (c.f. Pharmacological approach section of supplementary information). The pupae were briefly anesthetized on ice before injection and maintained on a cold aluminum block during the whole injection process. Each pupa received a microinjection of 66 nL dsRNA diluted in water which represents a concentration of 100 ng of dsRNA. The injected pupae were then placed in a plastic container of water (BioQuip®, Rancho Dominguez, CA, USA - 1425DG) to recover until emergence. The injection of 100 ng of *DOP1* dsRNA led to a survival of 50% of the pupae while 95% of the pupae emerged after being injected with the non-targeting control dsRNA (Fig. S6). The level of knockdown was assessed with RT-qPCR and Western blots (see supplementary information for details). We observed a decrease in the mRNA for *DOP1* in 60% of the injected mosquitoes and the knockdown was of about 30% (Fig. S6).

#### CRISPR/Cas9

The short guide RNAs (sgRNAs) used for CRISPR/Cas9 were designed to target the first exon of the conserved *DOP1* (AAEL003920). To define the sgRNA genomic target sites several factors were taken into account. Firstly, *Ae. aegypti* transcriptional databases were utilized to confirm RNA expression of putative target regions (*41*). We then performed blast searches to hunt for conservation and discovered an important conserved olfactory receptor domain termed 7tm-4 superfamily domain (pfam13853) that we decided to target (*42*). To minimize potential off-target effects, we confirmed specificity of our sgRNAs using publicly available bioinformatic tools (*43*) and selected the most specific sgRNAs within our target region. We produced these sgRNAs using *in vitro* transcription by combining primer pairs (primers 3 & 5) to make sgRNA-Target 1 and combining primers pairs (primers 4 & 5) to make sgRNA-Target 2. We then combined these sgRNAs (40 ng/µl) with purified Cas9 protein (300 ng/µl) purchased from PNA-bio (Newbury Park, CA, USA) and pre-blastoderm embryonic microinjections (*n*=300) were performed following previously established procedures (*35*). Following microinjection we individually isolated all surviving females (*n*=68), mated, blood fed, and allowed them to lay eggs. After egg laying, we isolated genomic DNA (Qiagen DNeasy Blood and Tissue Kit (Hilden, Germany)) from these females (focusing only on females that laid eggs (*n*=29)) and confirmed mutations in target sequences via PCR (standard techniques) with a primer pair that spans the cleavage sites amplifying 242bp of genomic DNA (primers 1 & 2). We discovered mutations in 68% (*n*=20/29) of the injected G0 females that laid eggs. We selected a mutant line (that stably transmitted the mutation to the G1 offspring) that generated an 18 nucleotide – 6 amino-acid deletion (LRRIGN) in the conserved 7tm-4 superfamily domain and backcrossed them, using individual female to male crosses every generation, for 9 generations. Mutations were verified using PCR/sequencing every generation (100% mutants for G5-G9). As additional controls, randomly selected mutant mosquitoes used in behavioural and electrophysiological assays were verified using PCR/sequencing after testing (100% were mutants), and electrophysiological AL recordings from *DOP1* mutants showed no significant changes in neuronal odour-evoked responses and spontaneous activity during dopamine superfusion (Fig. S11), verifying the efficacy of the CRISPR *DOP1* mutants. Primers and sgRNA sequences can be found in Supplementary Table S1.

#### Antibodies

The polyclonal antiserum against tyrosine hydroxylase (ImmunoStar, Hudson, WI, USA - Cat. no. 22941) was used at a concentration of 1:50 and monoclonal antisera against synapsin I (Sigma-Aldrich, St. Louis, MO, USA - Cat. No. WH0006853M7) were used at a concentration of 1:100 for immunohistochemistry. The antibody against the D1-like dopamine receptor, *DOP1* was custom made by 21st Century Biochemicals against a synthetic peptide corresponding to amino acids 138-154 of the *Ae. aegypti* protein, affinity purified, and used at a concentration of 1:100 for immunohistochemistry. This antibody was also used at a concentration of 1:1000 for western blot assays and recognizes a band with a mass of ∼72 kDa. Deglycosylation of protein samples with glycerol-free PNGase F (New England BioLabs, Ipswich, MA, USA - Cat. No. P0705) resulted in detection of a band at the expected molecular weight of ∼ 41 kDa. To further test specificity of this antibody, sections of *Ae. aegypti* brain tissue were divided into two wells and incubated with either antibody preadsorbed with 100 µM of the *DOP1* peptide (used to produce the antibody in rabbit) or with antibody alone and then processed for immunohistochemistry, as described below. Both wells were additionally incubated with antisera against synapsin I as a positive control for staining. Preadsorption with peptide from *DOP1* abolished *DOP1*-like immunoreactivity, while synapsin-like immunoreactivity remained (Fig. S6). Further details regarding immunohistochemistry and western blot assays are described in the supplementary information.

### Electrophysiology mosquito preparation

A total of 74 units recorded from 22 individuals, were exposed to a total of 418 odour stimulations in the electrophysiology experiments. Mosquitoes were immobilized on ice and mounted on a custom-designed holder (Fig. 4B) using UV-cured glue (Bondic®, Non Toxic Liquid Plastic Welder, BondicUSA, Fairfield NJ, USA). Each mosquito was tethered to the holder by the head capsule and the anterior-dorsal tip of the thorax, allowing steady electrophysiological recordings while the mosquito beats its wings in a fictive form of flight. All six legs were removed to prolong the flight bouts. A hole was cut in the cuticle of the head capsule to expose the antennal lobes, and then trachea and muscles 8 and 11 were removed. The brain was superfused continuously with temperature-controlled physiological saline solution (20° C) using a bipolar temperature controller and an in-line heater/cooler (CL-100 and SC-20, Warner Instruments) (Details on saline preparation and dopamine application are provided in the supplementary information).

### Coupled extracellular and behavioural recordings, spike sorting, and analysis

The tethered mosquito was placed on a Nikon FN-1 microscope (Eclipse FN1, Nikon Instruments Inc., Melville, NY, USA) under 20X objective (UMPlanFI, Olympus, Japan) to allow precise positioning of the recording electrode in one of the antennal lobes. Electrodes were pulled from quartz glass capillaries using a Sutter P-2000 laser puller and filled with 0.1 M LiCl. The electrode was positioned under visual control using the FN1 microscope and advanced slowly through the antennal lobe using a micromanipulator (PM10 - World Precision Instruments) until spikes were apparent in the recording channel. To determine the position of the recordings, the tip of each electrode was dipped into a solution of 2% Texas Red (ThermoFisher Scientific, Waltham, MA, USA) dissolved in 0.5 M potassium chloride solution before placement in the brain. After recording experiments, brains were imaged and z-stacks were taken at 1 µm steps using a two-photon microscope (Prairie Technologies Inc.).

Electrophysiological signals were amplified 10,000X and filtered (typically 0.1–5 kHz) (A-M Systems Model 1800, Sequim, WA, USA), recorded and digitized at 10 kHz using WinEDR software (Strathclyde Electrophysiology Software, Glasgow, UK) and a BNC-2090A analog-to-digital board (National Instruments, Austin, TX, USA) on a personal computer. Spike data were extracted from the recorded signal and sorted using a clustering algorithm based on the method of principal components (PCs) (Off-line Sorter; Plexon, Dallas, TX, USA). Only those clusters that were separated in three dimensional space (PC1–PC3) after statistical verification (multivariate ANOVA: p<0.1) were used for further analysis (2-6 units were isolated per preparation; n=22 preparations from as many mosquitoes; Fig. S7). Each spike in each cluster was time-stamped, and these data were used to create raster plots and to calculate peri-stimulus time histograms (PSTHs), interspike interval histograms, and rate histograms. All analyses were performed with R (R Core Team^45^) and Neuroexplorer (Nex Technologies, Winston-Salem, NC, USA) using a bin width of 20 ms, unless noted otherwise. We quantified the control corrected response for every unit by calculating a response index (RI). RI values reflect the deviation from the mean response of all units across all odors in one ensemble, as RI = (*R_odor_ - R_m_*)/*SD*, where *R_odor_* is the number of spikes evoked by the test odor minus the number evoked by the control stimulus, *R_m_* is the mean response, and *SD* is the standard deviation across the data matrix.

To couple electrophysiological and behavioural responses, we used a set-up (*44,46*) where an infrared camera (PointGrey Firefly MV FMVU-03MTC) was placed below the preparation. This set-up allowed an easy positioning of the recording electrodes, visualization of the flight responses, and stimulation of the preparation with olfactory stimuli (see supplementary information for details on *Olfactory Stimuli and Delivery*). IR LEDs were used to illuminate the wings, abdomen and proboscis, and images were recorded at 60 frames/s. A Python-based open source software (Kinefly (*47*)) calculated the wingbeat stroke amplitudes for each wing per frame. Because mosquito wing-beat frequencies are well above 400 Hz (and above the frame rate of the camera), we used a microphone (NR-23158-000, Knowles Electronics, LLC. Itasca, IL, USA), positioned below and adjacent to the preparation, to measure the wingbeat frequency. Wing stroke amplitude and wingbeat frequency were timestamped and acquired simultaneously with electrophysiological recordings.

## Acknowledgements

We thank B. Nguyen for mosquito colony maintenance; J. Joiner and K. Moosavi for assistance in olfactometer experiments; J. Stone for help with animal scent collections; and C. Bourgouin and M. Pereira for advice on the RNAi experiments. We thank P. Weir for comments and help with the arena experiments, and B. Brunton for statistical advice. We are grateful to D. Dickens for the scanned electron microscope images of *Ae. aegypti*. We acknowledge the support of the Air Force Office of Sponsored Research under grant FA9550-14-1-0398 and FA9550-16-1-0167, National Institutes of Health under grant NIH1RO1DCO13693-04, National Science Foundation under grant IOS-1354159, UC Riverside, MaxMind Inc., an Endowed Professorship for Excellence in Biology (JAR), the University of Washington Institute for Neuroengineering, and the Human Frontiers in Science Program under grant HFSP-RGP0022.

## Author Contributions

C.V., C.L., and J.A.R. conceived the study. C.V., C.L., participated in the execution and analysis of all aspects of the study. J.A.R. supervised and helped analyse the electrophysiology data presented in Figs. 4 and 5. G.H.W. generated and processed the immunohistochemistry data and western blots presented in Fig. 5 and S6. L.T.L. and J.E.L. helped carry out and analyse the behavioural assays presented in Figs. 1-4. J.Z.P. helped design the RNAi assays. O.S.A. designed and generated the CRISPR mutant mosquitoes. M.H.D. designed the flight arena experiments presented in Fig. 2. C.V., C.L. and J.A.R. wrote the paper, and all authors edited the manuscript.

